# Monocytes Strongly Induce (MYO)Fibroblast Contraction in a New 3D Skin Model to Understand the Inflammation-Fibrosis Axis in Systemic Sclerosis

**DOI:** 10.64898/2026.02.12.705496

**Authors:** D.C. Zanin-Silva, N.J.T. van Kooten, T.I. Papadimitriou, D.N. Dorst, B. Walgreen, E.L. Vitters, M.H.J van den Bosch, M.I. Koenders, A.P.M. van Caam

## Abstract

Systemic sclerosis (SSc) is an autoimmune disease characterized by excessive fibrosis and tissue stiffness, in which monocytes and macrophages are increasingly recognized as key contributors to pro-fibrotic myofibroblast formation and activation, although the underlying mechanisms remain incompletely understood. Here, we used a three-dimensional (3D) skin model to study how CD14^+^ monocytes, M1 and M2-like macrophages induce (myo)fibroblasts activation/contraction in collagen type I hydrogels. We identified that co-culture of fibroblasts with monocytes displayed strong spontaneous hydrogel contraction, coupled with an upregulation of myofibroblasts activation-associated markers, such as α-SMA and fibroblast activation protein. Using transcription-factor reporter constructs and small molecules inhibitors, we demonstrated that monocyte-fibroblast communication was mediated by JAK/STAT3 and TGF-β/Smad2/3 signaling pathways. Flow cytometry analyses revealed that monocytes, after interacting with fibroblasts, differentiated into a mixed M1/M2 polarization phenotype, characterized by CD163, CD206, CD86, and HLA-DR expression. Both M1 and M2-like macrophages promoted significant myofibroblast contraction, which could be mimicked by supernatant transfer. TGF-β neutralization but not IL-6 blocking abolished this effect. This study demonstrates that monocytes/macrophages can strongly induce (myo)fibroblasts activation/contraction. Together, our work contributes to elucidating pathways and mechanisms associated with skin fibrosis in SSc and paves the way for developing new platforms for targeted therapy testing.

## INTRODUCTION

Systemic sclerosis (SSc) is a severe autoimmune disease characterized by vasculopathy, immunological dysregulation, and fibrosis of the skin and internal organs^1^. The immune response plays a central role in the establishment and maintenance of SSc fibrosis, by promoting the differentiation, activation and survival of myofibroblasts^2^. Myofibroblasts are a subset of fibroblasts distinguished by their stress fibers, i.e. intracellular structures made of actin and myosin, that give them contractile properties, and their ability to produce high amounts of extracellular matrix (ECM) components, such as collagen type 1, type 3, and fibronectin. Notably, myofibroblasts express alpha-smooth muscle actin (α-SMA), which is associated with increased skin stiffness in SSc^3,4^.

Various immune cells, such as macrophages and their precursing monocytes, are implicated with myofibroblast activation during fibrotic conditions^5,6^. In SSc, monocyte/macrophage infiltration is seen across different affected organs, such as the lungs and the skin^7,8^. Recently, spatial transcriptomics on SSc patients’ skin biopsies revealed fibrotic niches enriched with fibroblasts and macrophages, closely linked to disease severity^9^. Moreover, biophysical changes in SSc circulating monocytes, including alterations in their cell area and deformation, reflect a pathological activation state and are associated with disease activity, microvascular damage, and fibrosis extent^10^. Nonetheless, the mechanisms by which monocytes/macrophages induce (myo)fibroblast activation and contraction in SSc remain incompletely understood.

Experimental models exploring mechanisms related to myofibroblast activation and contraction often incorporate key biomechanical properties of the fibrotic microenvironment in which these cells reside^11–13^. Collagen hydrogels present a valuable solution for this purpose, as they contain a component naturally produced by myofibroblasts, mimicking characteristics of a fibrotic ECM^14^. Within such 3D systems, cell co-cultures are essential to capture paracrine signaling and direct cell-cell interactions that regulate myofibroblast behavior^11,13^. However, few current SSc models incorporate immune cells into (myo)fibroblast cultures.

Here we used a three-dimensional (3D) skin model to understand whether monocytes/macrophages could cause myofibroblast activation and contraction in collagen type 1 hydrogels. We recently demonstrated a significant role for CD8^+^ T lymphocytes in inducing spontaneous (myo)fibroblasts contraction in these hydrogels^15^. However, it has not yet been elucidated whether other immune cells also implicated in the inflammation-fibrosis pathological axis in SSc, such as monocytes/macrophages, could exhibit a similar effect.

## RESULTS

### Monocytes strongly induce (myo)fibroblasts contraction in 3D collagen hydrogels

To investigate the potential role of monocytes in driving myofibroblast contraction in connective tissue diseases like SSc, we co-cultured healthy human dermal fibroblasts with CD14^+^ monocytes (isolated from peripheral blood mononuclear cells) in 3D collagen type 1 hydrogels.

Hydrogels containing only fibroblasts did not contract over 96 hours study (**Fig. 1a, b**), neither do hydrogels with only monocytes (data not shown). On the other hand, strong spontaneous contraction was observed in all hydrogels in which fibroblasts were co-cultured with monocytes, starting after 48 hours of the experiment (**Fig. 1b**). Co-culture of fibroblasts with PMBCs instead of monocytes, resulted in contraction of similar strength and comparable speed (**Fig. 1c**), equivalently to what we observed previously^15^. To test the relative contribution of monocytes to PBMCs-induced contraction, we depleted CD14^+^ cells before co-culturing with fibroblasts and observed that this significantly impaired hydrogels contraction (**Fig. 1d**), indicating that monocytes clearly promote myofibroblast contraction.

**Figure 1.**
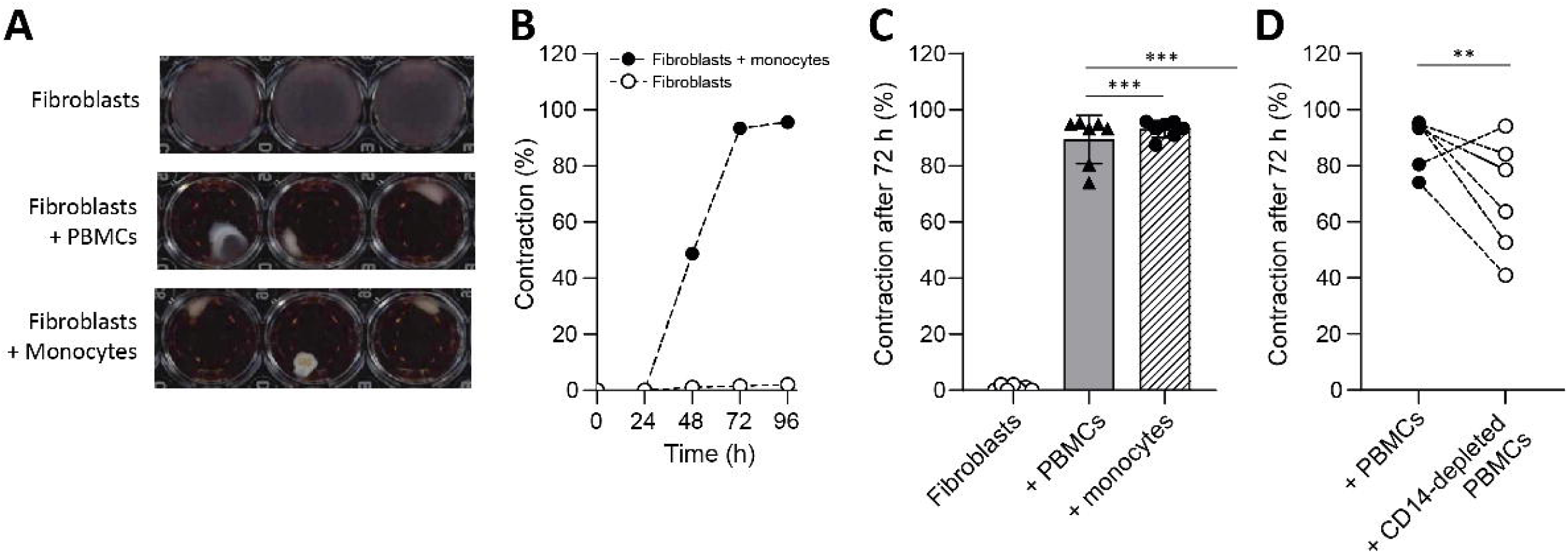
Monocytes strongly induce (myo)fibroblast contraction in the 3D skin collagen hydrogel model. (**A**) Representative scanned pictures of uncontracted hydrogels containing only fibroblasts or in co-culture with immune cells (PBMCs or monocytes). (Myo)fibroblast contraction is seen by a reduction in the surface area of the hydrogels. (**B**) Representative kinetics of (myo)fibroblast contraction over time (from one PBMC donor). Hydrogels with only fibroblasts did not contract. Inversely, hydrogels, in which fibroblasts were co-cultured with monocytes, displayed a robust contraction starting after 48 hours, reaching its maximum at 72 hours. (**C**) The average surface area of a contracted hydrogel group in triplicate divided by the total area of an uncontracted hydrogel. Co-culture of fibroblasts with monocytes or fibroblasts with PBMCs resulted in contraction of similar strength. (**D**) Depletion of CD14+ cells from PBMCs profoundly impaired hydrogels’ contraction. Each shape represents the average of 3 technical replicates of one PBMC donor, n = 7. Groups were statistically compared with one-way ANOVA with Tukey’s multiple comparisons test (**p<0.01, ***p<0.001).

### Monocytes lead to myofibroblasts activation markers expression

To study if and how monocytes also drive myofibroblast gene and protein expression, we co-cultured both cell types in 3D hydrogels and measured (pro-fibrotic) myofibroblast gene expression, such as α-SMA, FAP (fibroblast activation protein), and type 1 collagen^16,17^. To assess fibroblast specific gene expression, we used positive selection of CD45^+^ cells, as fibroblasts are negative for this CD marker. 24 hours after co-culture with monocytes, fibroblasts exhibited a significant increase in gene expression of *PDPN* (podoplanin), *COL3A1*, *HLA-DR*, *PLOD2* (Procollagen-Lysine,2-Oxoglutarate 5-Dioxygenase 2), and *IL-6* (interleukin-6) (**Fig. 2a**), reflecting an activated and pro-fibrotic status.

**Figure 2.**
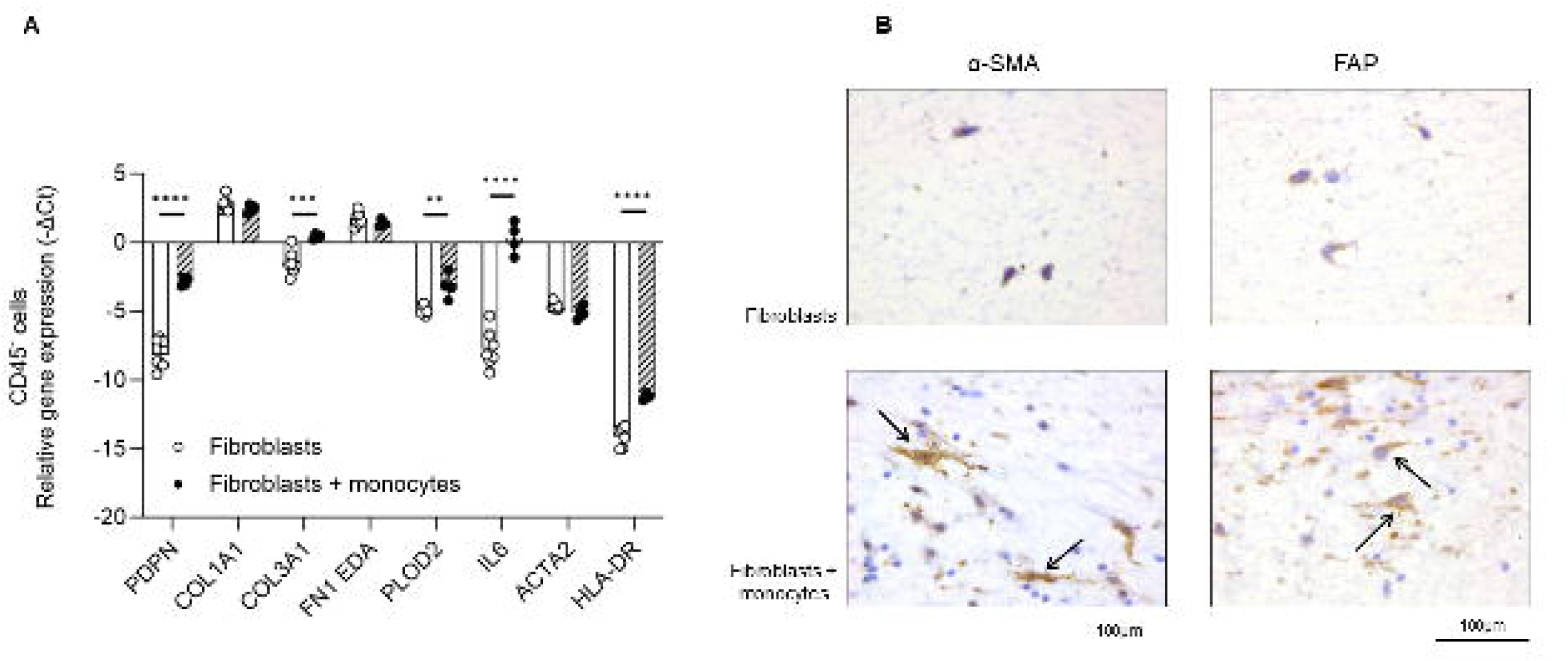
Monocytes promote robust expression of different myofibroblast activation markers. (**A**) Sorted CD45-cells (fibroblasts) were analyzed for expression of myofibroblast-related genes after co-culture with monocytes in collagen hydrogels. Values (mean ± SEM) indicate relative gene expression (−ΔCt) measured with qPCR. Reference genes used were GAPDH and RPS27A. Analyses were done by two-way ANOVA with Sidak’s multiple comparisons test (**p<0.01, ***p<0.001, ****p<0.0001). (**B**) Representative immunohistochemistry staining showing robust expression of α-SMA (alpha-smooth muscle actin) (left) and fibroblast activation protein (FAP) (right) in the hydrogels containing only fibroblasts (top) compared to those co-culture with monocytes (bottom). The small blue nuclei in the bottom panels are monocytes. (Myo)fibroblasts are clearly positive for both markers (black arrows), with a typically elongated shape.

Subsequently, we confirmed this monocyte-induced myofibroblast phenotype at protein level through immunohistochemical staining of histological sections of the skin hydrogels. Fibroblasts co-cultured with monocytes were clearly positive (indicated by arrows) for fibroblast activation protein (FAP) and alpha-smooth muscle actin (α-SMA), indicating strong fibroblast activation by monocytes (**Fig. 2b**).

### Monocyte-induced JAK/STAT and TGF-β/Smad signaling pathways in dermal fibroblasts during co-culture

To elucidate the mechanisms underlying myofibroblast activation, we analyzed which intracellular signaling pathways are triggered in fibroblasts in response to monocytes. For this, we used dermal fibroblasts previously transfected with transcription factor reporter constructs^18^, including Smad-binding elements (SBE) to capture canonical TGF-β signaling, STAT-inducible elements (SIE) reflecting JAK/STAT activation, NF-κB response elements representing inflammatory signaling, and interferon-stimulated response elements (ISRE) indicative of type I interferon responses.

After 24 hours of co-culture, we observed in the fibroblasts that NF-κB (**Fig. 3a**) and SIE (**Fig. 3b**) constructs exhibited strong activity in the presence of monocytes. Inversely, reduced activity was found using SBE (**Fig. 3c**) and ISRE (**Supp Fig. 1a**) constructs. No changes were found in the responses of NFAT-5 or CRE constructs after co-culture with monocytes (**Supp Fig. 1b, c**), indicating that pathways associated with osmotic stress responses and cAMP/PKA signaling were not involved.

**Figure 3.**
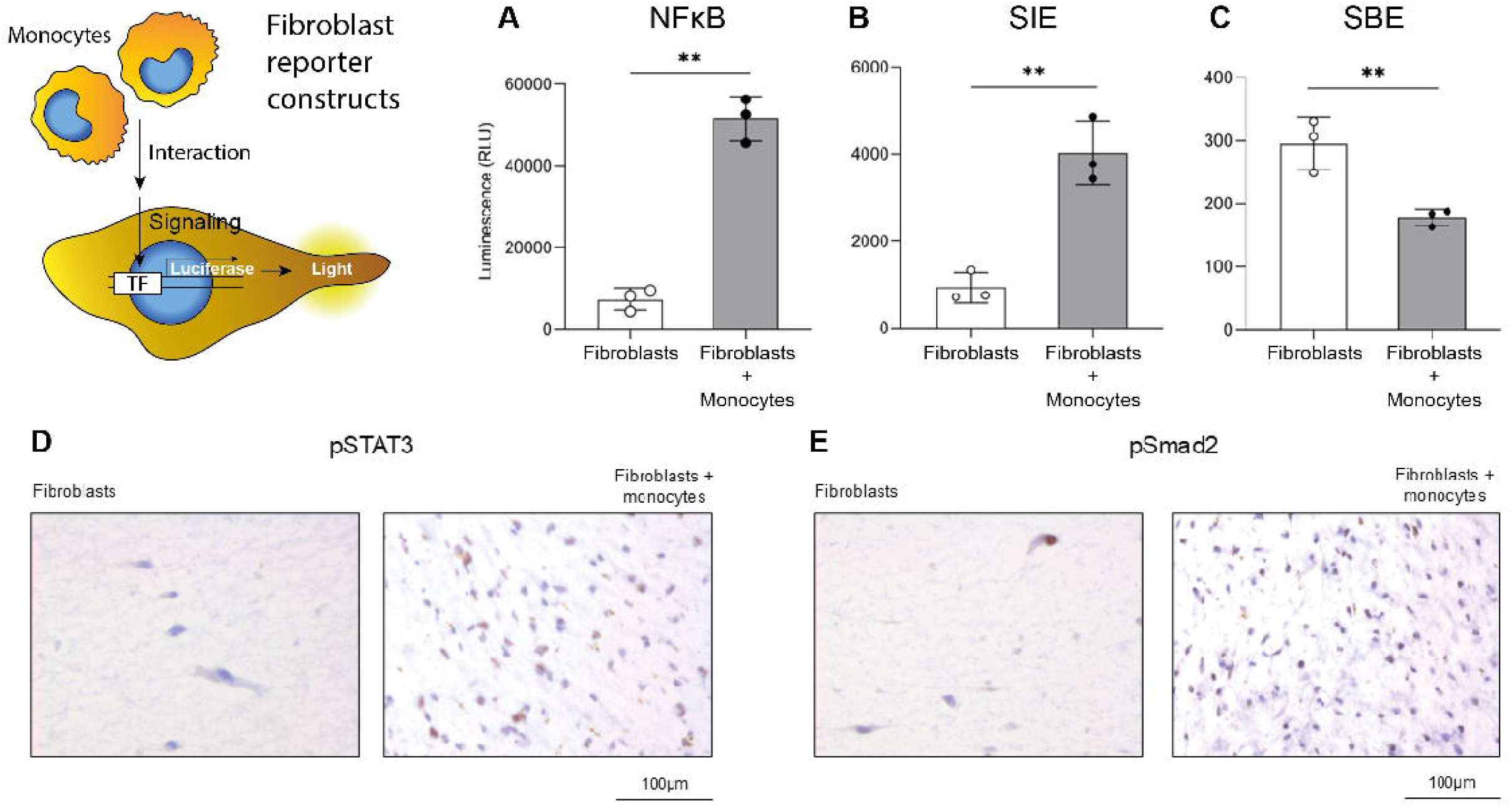
Signaling pathways activated in fibroblasts constructs during the co-culture with monocytes. Dermal fibroblasts, expressing transcription factor-responsive reporter constructs, were used to monitor the activation of relevant signaling pathways. Following stimulation (by co-culture with monocytes), transcriptional activation drives luciferase expression, which is subsequently quantified by luminescence as an indirect measure of pathway activity. Luminescence values were measured in fibroblasts with NF-κB (**A**), SIE (**B**), and SBE (**C**) reporter constructs in the presence of monocytes (for 24 hours) in the collagen hydrogels. Hydrogels with only fibroblasts (without any stimulation) were used as negative controls. Statistical analyses were performed using Student’s t-test (**p<0.01). Each dot represents the average of 3 technical replicates of one PBMC donor, n = 3. The phosphorylation of STAT3 (pSTAT3) (**D**) and Smad2 (pSmad2) (**E**) was clearly observed by immunohistochemistry in monocyte-containing hydrogels (right) when compared to fibroblasts cultured alone (left), demonstrating induction of TGF-β and STAT3 signaling in the co-culture. NF-κB: Nuclear factor-κB response element; SIE: Interleukin (IL)-6 sis-inducible element or STAT3 response element, and SBE: SMAD binding element.

The induction of pathways associated with pro-inflammatory response of fibroblasts, such as SIE, was subsequently confirmed by immunohistochemistry through clearly observed phosphorylation of STAT3 (pSTAT3) in the hydrogels (**Fig. 3d**). In the hydrogels containing only fibroblasts, pSTAT3 staining was not detected (**Fig. 3d**). Additionally, the unexpected results showing reduced SBE (Smad binding element) activity (**Fig. 3c**) led us to further investigate the activation state of other components associated with this pathway. In contrast to the reduced SBE reporter signal, immunohistochemistry analysis revealed strong Smad2 phosphorylation in fibroblast during the co-culture with monocytes (**Fig. 3e**), indicating that receptor-proximal Smad activation does not directly translate into increased gene expression due to more downstream regulatory processes^19^. Together, these results suggest that the JAK/STAT and TGF-β/Smad pathways orchestrate monocyte-induced myofibroblast activation in our 3D co-culture model.

### Essential role for JAK/STAT and TGF-β/Smad signaling in (myo)fibroblasts contraction induced by monocytes

To determine whether the activation of the observed pathways was also involved with myofibroblast contraction, we treated the co-culture hydrogels with different pharmacological inhibitor. For this, we used tofacitinib, a JAK/STAT signaling inhibitor^20^ and/or SB-505124, a selective inhibitor targeting the type 1 TGF-β receptor (TGFBR1) and thus Smad signaling^21^. We also tested two other compounds known to block relevant signaling pathways in fibroblasts: IKK-16, a selective inhibitor of IκB kinase (IKK)^22^, and TAK-242, a toll-like receptor 4 (TLR4) signaling inhibitor^23^.

3D hydrogels containing fibroblasts plus monocytes treated from the start with SB-505124 exhibited a significant reduction in contraction compared to those treated with only vehicle DMSO (**Fig. 4**). When SB-505124 was added to the culture medium one day (24 hours) after the hydrogels’ preparation, no inhibition of contraction was observed (**Fig. 4**), suggesting that early activation of TGF-β signaling is a critical step in the model. In the hydrogels receiving IKK-16 or tofacitinib (**Fig. 4**), no blockade of contraction was observed, indicating that although monocytes induce these signaling pathways in fibroblasts, they are not involved in contraction. Interestingly, tofacitinib more strongly inhibited hydrogel contraction if it was added after 24 hours instead of at the start of the experiments (**Fig. 4**), which indicate that JAK/STAT signaling is activated or only relevant at a later stage. Blocking the TLR4 signaling did not affect (myo)fibroblasts contraction (**Supp Fig. 2**), indicating that damage-associated molecular pattern (DAMP) signaling is not involved with myofibroblast contractile activity in this context.

**Figure 4.**
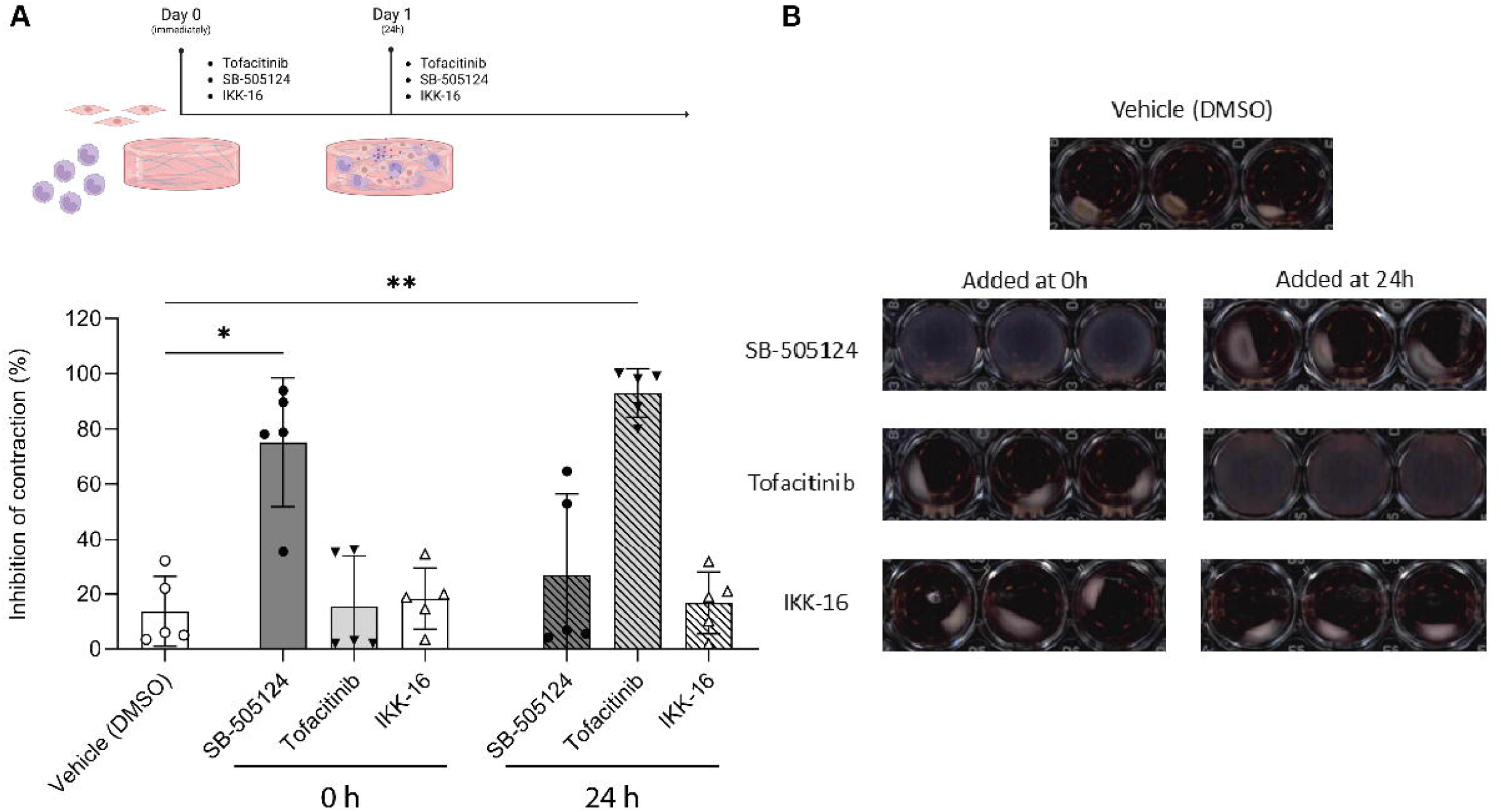
TGF-β/Smad and JAK/STAT signaling pathways are important regulators in monocyte-driven (myo)fibroblast contraction. The co-culture of fibroblasts and monocytes was treated with SB-505124 (5 µM), tofacitinib (1 µM), or IKK-16 (1,5 mM) either immediately (0h) after hydrogels generation or 24h later and subsequent contraction at 72h was compared with vehicle-treated (DSMO) (control). (**A**) Hydrogels treated immediately (0h) with SB-505124 displayed impaired contraction, which did not occur when the inhibitor was added later (at 24h). Inversely, hydrogels receiving tofacitinib at 0h showed strong spontaneous contraction, which did not happen upon treatment after 24h. Blocking the NF-κB pathway did not affect (myo)fibroblast contraction at any time point. Each shape represents the average of 3 technical replicates of one PBMC donor, n = 5. Groups were statistically compared with one-way ANOVA followed by Tukey’s multiple comparisons test (*p<0.05, **p<0.01). (**B**) Representative scanned images of the co-cultured hydrogels treated with SB-505124, tofacitinib or IKK-16 at the different mentioned time points (0h or 24h).

Summarily, the use of inhibitors allowed us to identify JAK/STAT and TGF-β/Smad as key signaling pathways involved in monocyte-induced (myo)fibroblast contraction.

### Interaction with fibroblasts leads monocytes to differentiate into macrophages with a mixed M1/M2-like phenotype

We further examined how direct interaction with the fibroblasts within the hydrogels shaped monocyte differentiation. Once in the tissue, monocytes can differentiate into macrophages, and CD68, a type I transmembrane glycosylated protein, is the molecule commonly used as a marker to identify these cells in humans^24,25^. Examining the histology of our co-cultured hydrogels, we identified clear CD68 expression on day 3 of the experiment, suggesting monocyte-to-macrophage differentiation (**Fig. 5a**). Interestingly, we noted that many of the CD68^+^ cells were distributed forming a sort of “surrounding layer” covering the fibroblasts in the hydrogels, a phenomenon also seen during wound healing processes^26,27^.

**Figure 5.**
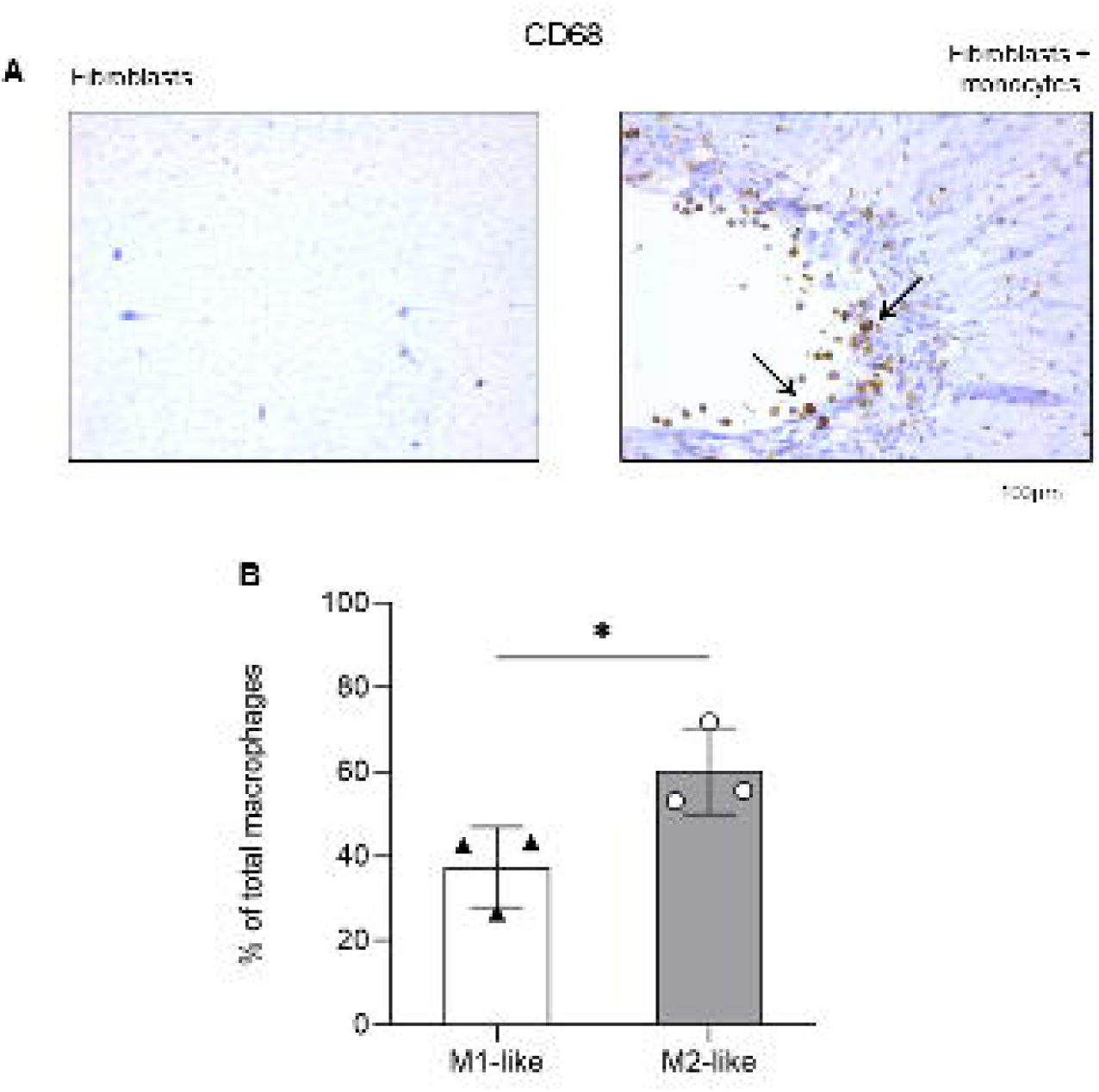
Interaction with fibroblasts promotes monocyte-to-macrophage differentiation. (**A**) Representative immunohistochemistry staining showing clear CD68 expression in the monocyte-containing hydrogels (right). Black arrows indicate a “surrounding layer” formed by the CD68^+^ cells covering the fibroblasts. (**B**) Flow cytometry analyses showing the percentage of total macrophages (CD45^+^CD68^+^) isolated from the hydrogels after enzymatic digestion and subsequently CD45^+^ magnetic selection. Cells (CD45^+^CD68^+^) were classified into M1-like (CD163^−^) or M2-like (CD163^+^) macrophages. Comparisons were performed using Student’s t-test (*p<0.05). Each shape represents the average of 3 technical replicates of 1 PBMC donor, n = 3.

After enzymatic digestion of the hydrogels, the CD45^+^ cell fraction was selected and stained for flow cytometry analysis to assess their immunophenotypic profile. We identified macrophages based on the expression of CD68 (CD45^+^CD68^+^) and subsequently classified them as M1– or M2-like macrophages (the gating strategy is detailed in **Supp Fig. 3**) based on the absence or presence of scavenger receptor CD163 respectively^28,29^.

Our analyses demonstrated that co-culture with fibroblasts led to the appearance of a significant percentage of M2-like macrophages (CD45^+^CD68^+^CD163^+^) (**Fig. 5b**) compared to M1-like cells (CD45^+^CD68^+^CD163^−^). As expected, M2-like macrophages were positive for the expression of CD206 (mannose receptor) (**Supp Fig. 3**), a classical marker of M2 profile^30,31^. Moreover, we observed that M1-like macrophages, in addition to the expression of markers traditionally associated with their activation state, such as CD86 and HLA-DR^29^, also expressed CD206 (**Fig. 4a**). In a similar manner, M2-like cells expressed markers associated with M1 polarization (**Fig. 4b**), HLA-DR and CD86, suggesting that interaction with fibroblasts induces monocytes to differentiate into macrophages with a mixed M1/M2 polarization phenotype, which may reflect a contribution to both proinflammatory and fibrotic processes.

### In vitro-generated M1– and M2-like macrophages promote (myo)fibroblasts activation and contraction

Finally, we aimed to verify whether polarized M1 or M2 macrophages could also induce (myo)fibroblasts activation and contraction. Although classically activated M1 macrophages are known for their pro-inflammatory role, they are also described as important players during fibrotic conditions, including in the lungs^32^ and peritoneum^33^. On the other hand, M2 macrophages are primarily associated with tissue repair and regeneration and are more commonly linked to fibrosis^34,35^. To verify each cell subset influence, we produced hydrogels containing fibroblasts and macrophages derived from monocytes, which were polarized in vitro using a cytokine cocktail (see Methods) into the two well-known activation states: GM-CSF-generated M1 or M-CSF-generated M2. We observed that two days (48h) after hydrogels’ preparation, both M1– and M2-like cells were strongly capable of inducing (myo)fibroblasts contraction, as seen by a reduction in hydrogels’ area (**Fig. 6a**). To assess whether this effect could be attributed to released soluble mediators, we used the culture supernatant from M1– and M2-like macrophages to stimulate collagen hydrogels containing only fibroblasts. Although hydrogels receiving either M1 or M2-like supernatant both exhibited strong spontaneous contraction after four days (96h) of culture (**Fig. 6a**), this response was less pronounced than with direct co-culture, suggesting that cell-cell contact plays a significant role in promoting myofibroblast contraction.

**Figure 6.**
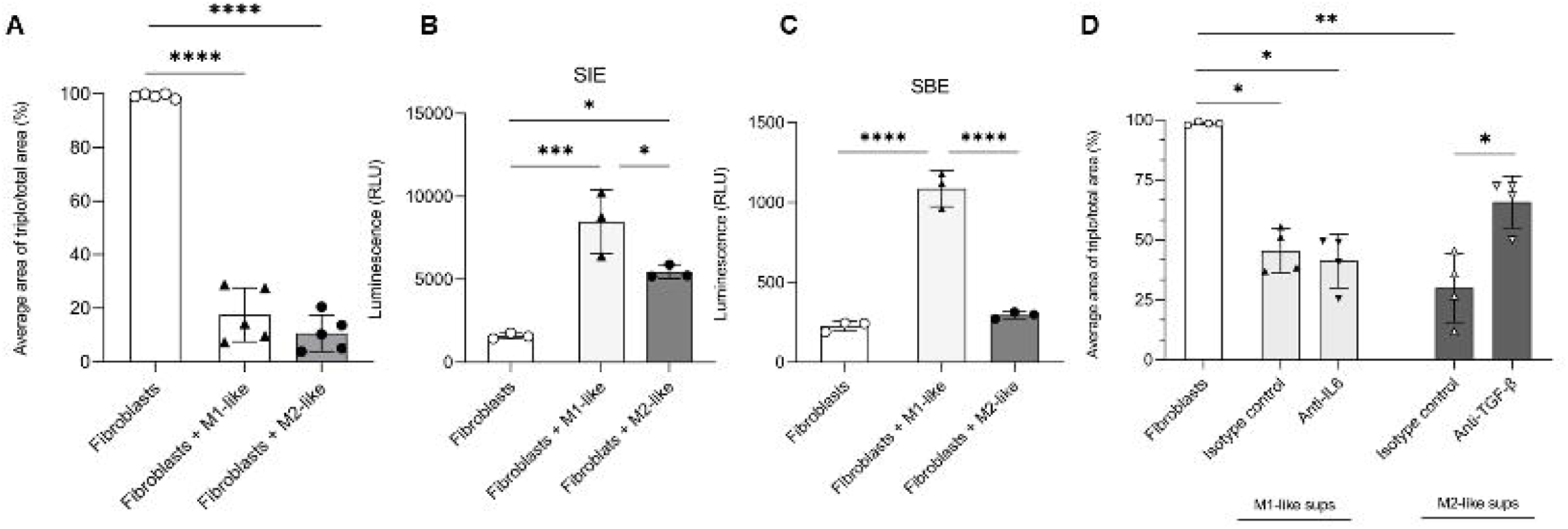
M2-like macrophages induce (myo)fibroblast contraction through TGF-β in the hydrogels. (**A**) Hydrogels containing co-culture of fibroblasts and in vitro-generated M1– or M2-like macrophages showed a strong spontaneous contraction, reaching its maximum at 48 hours (not shown). Hydrogels with only fibroblasts did not contract. Each shape represents the average of 3 technical replicates of one PBMCs donor, n = 5. Groups were statistically compared with one-way ANOVA and Tukey’s multiple comparisons test (****p<0.0001). (**B**, **C**) To assess signaling pathway activation, luminescence values measured in fibroblasts transfected with SIE (**B**) and SBE (**C**) reporter constructs in the presence (for 24 hours) of M1– or M2-like macrophages in the collagen hydrogels. Hydrogels with only fibroblasts were used as negative controls. Statistical analyses were performed using Student’s t-test (*p<0.05, ***p<0.001, ****p<0.0001). Each shape represents the average of 3 technical replicates of 1 PBMC donor, n = 3. (**D**) M1 or M2-like macrophage culture supernatants were incubated for 1h with neutralizing antibodies against IL-6 or TGF-β, respectively. An IgG isotype antibody was added as a control. Blocking TGF-β significantly impaired M2-like supernatant-induced contraction. Differently, IL-6 neutralization in the M1-like supernatant had no effect on preventing (myo)fibroblasts contraction. Each shape represents the average of 3 technical replicates of 1 PBMC donor, n = 4. Groups were statistically compared using one-way ANOVA with Tukey’s multiple comparisons test (*p<0.05, **p<0.01).

Using the reporter fibroblasts constructs once more, we observed that M1-like macrophages induced significant SIE activity, more than M2-like cells (**Figure 6b**). Similarly, fibroblasts also exhibited increased SBE activity in the presence of M1 (**Fig. 6c**) compared to M2-like cells. Comparable results were seen for NF-kB and ISRE reporters (**Supp Fig. 5a, b**), whereas no changes were detected in CRE and NFAT-5 cells (**Supp Fig. 5c, d**), suggesting that M1-like macrophages have a greater capacity to stimulate inflammatory signaling in fibroblasts.

Finally, we aimed to link the results of the enhanced SIE (JAK/STAT) pathway and SBE (SMAD) pathway to potential soluble mediators: IL-6 and TGF-β. For this, we incubated M1-like supernatants with anti-IL-6 and M2-like supernatants with anti-TGF-β neutralizing antibodies, based on the predominant cytokine profile of each macrophage subtype, and then analyzed if these cytokines mediated (myo)fibroblast contraction. Hydrogels treated with M2-like supernatant incubated with TGF-β neutralizing antibody displayed a reduction in the contraction process compared to those treated with an isotype control antibody (**Fig. 6d**), supporting a role for M2-like cell-derived TGF-β in driving (myo)fibroblast contraction. On the other hand, IL-6 neutralization in the M1-like supernatant had no effect on preventing (myo)fibroblast contraction in the hydrogels (**Fig. 6d**), pointing to the involvement of additional M1-like cell-derived mediators in this process.

## DISCUSSION

During homeostasis, myofibroblasts play an essential role in tissue remodeling and the wound healing process^36^. However, their sustained activation can substantially contribute to the development of various fibrotic diseases, such as systemic sclerosis^4^ and Dupuytren’s contracture^37^. Here, we employed a three-dimensional (3D) skin collagen type 1 hydrogel model consisting of a co-culture of healthy primary monocytes/macrophages and dermal fibroblasts, enabling the assessment of immune cell-driven (myo)fibroblast activation and contraction, which reflects increased skin stiffness and fibrosis in systemic sclerosis (SSc). Through the co-culture, we observed a crucial role of monocytes in these processes. Notably, enhanced (myo)fibroblasts contractility was accompanied by upregulation of certain genes such as PDPN, COL3A1, HLA-DR, PLOD2, and IL-6, reflecting an activated and pro-fibrotic myofibroblast phenotype. These genes have been also found elevated in SSc patients’ skin myofibroblasts, coupled with increased levels of α-SMA, FAP, and type 1 collagen^38–41^, suggesting that, in our model, monocytes drive fibroblasts to adopt a disease-relevant phenotype.

Further supporting this, we identified at protein level (myo)fibroblasts strongly positive for α-SMA and FAP expression following co-culture with monocytes. Notably, dermal fibroblasts isolated from SSc patients’ skin, when embedded in a 3D microenvironment such as biocompatible polyisocyanide (PIC-RGD) hydrogels, retain critical features of their in vivo behavior, including increased contraction, proliferation, and expression of α-SMA, collagen, and FAP. Inversely, under conventional 2D culture conditions, these cells rapidly lose their pro-fibrotic phenotype^42^. In this context, our results reinforce the importance of a 3D microenvironment not only in maintaining the pro-fibrotic myofibroblasts features induced by monocytes but also in enabling functional crosstalk between immune and stromal cells, which is largely absent or blunted in traditional 2D culture systems. Thus, the monocyte-induced upregulation of α-SMA and FAP reflects a biologically relevant activation of myofibroblasts, occurring in a setting that more closely mimics the in vivo fibrotic niche.

Recently, the interaction between macrophages and fibroblasts in SSc has gained more attention. An elegant spatial multi-omics study integrating data from SSc patients and mouse models identified the co-localization and signaling dynamics of fibroblasts and macrophages within fibrotic skin niches^9^. This disease-specific microenvironment enriched in both cell types was characterized by elevated TGF-β pathway activity and a strong correlation with patients’ clinical outcomes. The results suggest that fibrotic progression is an intricate process involving both the close spatial proximity of immune and stromal cells and dynamic signaling interactions occurring within organized niches. Sequential events, such as immune cell infiltration and disrupted cell communication, might progressively destabilize tissue homeostasis, promoting a shift toward a pro-fibrotic state^9^. In this context, our work adds a functional perspective by showing how infiltrating cells such as monocytes directly reshape fibroblast behavior, highlighting the phenotypic and mechanical transformation of fibroblasts into contractile myofibroblasts. We identified that monocyte-driven (myo)fibroblast activation in the hydrogels involved signaling pathways that are critical in SSc, such as the IL-6/STAT3 and TGF-β/Smad pathways. In SSc patients’ skin, dermal fibroblasts and infiltrating mononuclear cells exhibit a prominent IL-6 expression. Additionally, elevated IL-6 levels are associated with more extensive skin involvement and reduced long-term survival^43^. Upon binding to its receptor (IL-6R), IL-6 can activate the JAK/STAT pathway, leading to STAT3 phosphorylation^44^. STAT3 is then translocated to the nucleus, binding to SIE, which consists of repeats of “IL-6 sis-inducible elements,” also known as STAT3 response elements, resulting in the expression of genes involved in cell proliferation, inflammatory response, and fibrogenesis^45,46^. Here, we demonstrated a clear phosphorylation of STAT3 in the hydrogels containing monocytes and fibroblasts. In addition, SIE-reporter fibroblasts activity significantly increased in the presence of monocytes, indicating that the JAK/STAT3 pathway plays a key role in monocyte-induced (myo)fibroblasts activation.

While IL-6 blocking therapies in SSc, such as tocilizumab, have demonstrated only modest effects in cutaneous fibrosis^47^, growing evidence suggests that targeting downstream signaling such as JAK/STAT3 could provide valuable clinical outcomes^48,49^. Therefore, our findings support a role for STAT3 in monocyte-induced myofibroblast activation and strengthen the argument that targeting downstream pathways such as JAK/STAT3 may offer an effective approach in diseases like SSc with multifactorial cytokine involvement.

TGF-β/Smad signaling pathway is another important regulator of fibrosis, directly influencing the activation and differentiation of myofibroblasts^50^. Multiple abnormalities in the Smad signaling pathway have been detected in SSc^51^, including elevated phosphorylation of Smad2/3 in cultured SSc patients dermal fibroblasts^52,53^. In our study, we detected that monocytes led to Smad2 phosphorylation when co-cultured with fibroblasts in the hydrogels, further supporting the acquisition of an activated and pro-fibrotic phenotype. However, unexpectedly, SBE (Smad Binding Elements)-reporter fibroblast constructs exhibited decreased activity in the presence of monocytes. A reduction of Smad3 nuclear translocation was detected in myofibroblasts isolated from a murine model of scleroderma, while Smad2 translocation showed no changes^19^, indicating that some pro-fibrotic actions of TGF-β occur through distinct modulation of Smad2 and Smad3 activities. These differences in transcriptional activation mechanisms by Smad2 and Smad3^54^, as well as potential post-translational modifications, could explain our contrasting results between increased Smad2 phosphorylation in the hydrogels and decreased activity of SBE-reporter fibroblasts after co-culture with monocytes.

Building on these mechanistic insights, we also explored whether targeting the identified signaling pathways could modulate (myo)fibroblasts contraction in our 3D model. Treatment with SB-505124, a TGF-β receptor type 1 inhibitor, at the beginning of the co-culture reduced drastically contraction on hydrogels containing fibroblasts and monocytes. In contrast, a later treatment (24 hours after co-culture initiation) had no significant effect, suggesting that early activation of the TGF-β/Smad signaling pathway is a crucial step for myofibroblast contraction. SB-505124 has previously been shown to reduce collagen deposition and myofibroblast numbers in an experimental model of fibrosis^55^, highlighting our 3D co-culture hydrogels as a relevant platform for testing therapies within this context. On the other hand, in an unexpected manner, immediate treatment of co-culture hydrogels with tofacitinib did not reverse (myo)fibroblasts contraction, however, it completely blocked this process when added later. Tofacitinib is a JAK1/JAK3 pathway inhibitor that has demonstrated potential efficacy in treating SSc patients, reducing skin thickness and leading to positive impacts on musculoskeletal involvement^56^. Here, we believe that as co-culture progresses, pro-inflammatory cytokines, possibly secreted by monocytes and/or (myo)fibroblasts, could induce JAK/STAT pathway, explaining why tofacitinib had a stronger effect when added after one day of experimentation.

Having established that monocytes drive (myo)fibroblast activation and contraction in our 3D skin model, we next investigated how monocytes themselves are modulated through interaction with fibroblasts, aiming to capture the bidirectional nature of these cells’ crosstalk. The co-culture of monocytes and fibroblasts in the hydrogels led to a distinct pattern of CD68 expression, with numerous CD68⁺ cells forming a surrounding layer covering the fibroblasts. This spatial arrangement bears some resemblances observed during wound healing process, where fibroblast-mediated signals guide macrophage positioning to support tissue repair^27^. Moreover, deformation fields generated by fibroblasts in fibrillar collagen matrices have been shown to dynamically attract macrophages over distances, coordinating their migration and activation^26^. Such similarities could indicate that our co-culture model recapitulates key aspects of the physiological crosstalk between fibroblasts and immune cells, supporting its biological relevance.

We further identified and classified the CD68^+^ cells into M1 or M2-like macrophages based on the expression of the scavenger receptor CD163. Flow cytometry analyses revealed a higher percentage of M2-like macrophages (CD45^+^CD68^+^CD163^+^) in the hydrogels following the co-culture with the fibroblasts. Consistent findings were reported in a human skin equivalent model, co-culturing SSc patients’ monocytes and fibroblasts, in which monocytes differentiated into CD163⁺ and CD206⁺ macrophages^57^. Here, many of the M2-like cells were found also expressing markers commonly associated with M1 macrophages, such as HLA-DR and CD86. Similarly, M1-like macrophages (CD45^+^CD68^+^CD163^−^) exhibited markers related to the M2 phenotype, such as CD206, indicating the occurrence of a mixed M1/M2 polarization phenotype. Such mixed phenotype was observed in circulating monocytes from SSc patients^58,59^ and has also been reported in other pathological contexts, including lung cancer^60^ and osteoarthritis^29^. Our findings suggest that cells’ interaction within the hydrogels might lead to the release of inflammatory and pro-fibrotic mediators, fostering a spectrum of macrophage polarization state rather than a strict M1/M2 dichotomy.

Although M2 macrophages are widely associated with fibrosis, their isolated contribution may represent an oversimplified perspective^5^. In fact, a study using a murine model of SSc has shown that blocking M1 macrophage polarization can serve as an effective therapeutic strategy, particularly in the early stages of the disease^61^. These findings indicate a potential involvement of M1 pro-inflammatory macrophages in triggering or amplifying fibrotic responses, while M2-like macrophages contribute to sustaining and maintaining fibrosis over time. Here, when co-culturing fibroblasts with M1 or M2-like cells in the 3D collagen hydrogels, we detected strong spontaneous contraction of (myo)fibroblasts in the presence of both macrophage subtypes, an effect similarly observed using M1 and M2-like culture supernatant to stimulate hydrogels containing only fibroblasts, emphasizing that both M1– and M2-like macrophages contribute to myofibroblast activation. Notably, the faster contraction observed in the co-culture condition (within 2 days), compared to delayed effects seen with supernatant exposure alone (after 4 days), points to a significant role for direct cell–cell contact in accelerating the fibrotic response^62^. Lastly, we pre-incubated the respective macrophage supernatants with neutralizing anti-TGF-β or anti-IL-6 antibodies to investigate the contribution of released soluble factors. We observed a marked reduction in (myo)fibroblast contraction when M2-like supernatants were treated with anti-TGF-β, whereas neutralizing IL-6 in M1-like supernatants did not yield the same effect. These results reinforce the crucial role of TGF-β in (myo)fibroblast contraction and suggest that additional mediators released by M1-like macrophages, such as IL-1 or reactive oxygen species (ROS), might also contribute to the contraction process.

Despite providing valuable insights, our study has certain limitations. Incorporating only fibroblasts and monocytes/macrophages does not fully recapitulate the complexity of the pathological triad in SSc. Therefore, adding endothelial cells into 3D skin co-culture systems^63,64^ could provide a more comprehensive alternative for studying SSc-related vasculopathy, a crucial parameter in the disease pathogenesis and patients’ clinical outcomes^65^. Finally, the limited availability of isolated SSc patient cells (monocytes/macrophages and fibroblasts) prevented their inclusion in our study. Future research will focus on developing a personalized in vitro skin model by combining donor cells, with experiments involving SSc patient samples.

## METHODS

### Isolation and culture of primary skin dermal fibroblasts

Human primary dermal fibroblasts were obtained from punch skin biopsies (4 mm) of the dorsal mid-forearm of a healthy male donor under local asepsis and anesthesia. The procedure and samples collection were done after written informed consent form (study number: NL57997.091.16). The material was placed in a 24-wells plate containing 2 ml DMEM Glutamax medium (Gibco, USA), supplemented with 100 U/ml penicillin, 100 mg/ml streptomycin, 100 mg/L pyruvate, and 20% fetal calf serum (FCS). Plates were incubated in appropriate culture conditions (37 °C, 5% CO2, 95% humidity) for 2 weeks until the spontaneous outgrowth of the primary fibroblasts. Every 3-4 days, the medium was partly refreshed. Subsequently, cells were cultured and expanded on plastic in T175 flasks in DMEM Glutamax medium supplied with 100 U/ml penicillin, 100 mg/ml streptomycin, 100 mg/L pyruvate, and 10% FCS. Fibroblasts were used up to a maximum number of 12 passages.

### Human monocyte isolation and differentiation to M1 or M2-like macrophages

Healthy monocytes were isolated from human peripheral blood mononuclear cells (PBMC) obtained from buffy coats (project number: NVT 0397-02) supplied by Sanquin blood bank (Nijmegen, The Netherlands) by Ficoll^®^-Paque PLUS (Sigma-Aldrich, USA) density centrifugation, following manufacturer’s guidelines. PBMC passed through a magnetic-activated cell sorting (MACS) isolation, using CD14^+^ magnetic MicroBeads (130-050-201, Miltenyi Biotec, Germany) and MACS buffer: Phosphate Buffered Saline (PBS) with 1% FCS and 1.5% Anticoagulant Citrate Dextrose Solution (ACD)-A.

For macrophage differentiation, as previously described^29^, the obtained monocytes were seeded in 48-wells Upcell^TM^ plates (Thermo Fisher Scientific, USA) at 250,000 cells/well and cultured in RPMI 1640 GlutaMAX^TM^ medium (Gibco) supplemented with 10% FCS, 100 mg/L pyruvate, 100 U/mL penicillin, and 100 mg/mL streptomycin. Cells were differentiated during 6 days to M1-like macrophages using 50 ng/mL granulocyte-macrophage colony-stimulating factor (GM-CSF) (7954-GM-050/CF, R&D System, USA) or to M2-like macrophages using 20 ng/mL macrophage colony-stimulating factor (M-CSF) (216-MCC-025/CF, R&D Systems). On day 3 of culture, the medium with supplements was refreshed. Cells were activated on day 5, by adding 20 ng/mL interferon (IFN)-γ (570202, BioLegend, USA) or 10 ng/mL interleukin (IL)-4 (204-IL, R&D Systems) for the M1 or M2-like macrophages, respectively. The supernatant was collected on the last day of culture (day 6), spun at 400×g for 10 min to remove cell debris, and stored at –80 °C until use. Finally, cells were detached and collected by 1-hour incubation in PBS at 4°C.

### 3D skin co-culture of monocytes/macrophages and fibroblasts in a collagen hydrogel contraction model

Primary skin dermal fibroblasts were washed with saline and detached from the culture flasks using trypsin (T1426-100MG, Sigma-Aldrich, USA) at 37°C. Subsequently, 10 ml of DMEM medium with 10% FCS was added to inactivate the trypsin. PBMC, isolated monocytes (CD14^+^), PBMC without monocytes (depleted from CD14^+^ cells by MACS), and M1 or M2-like macrophages were obtained as described before. To make each 3D collagen hydrogel^15^, 20 µl 10 x Minimum Essential Medium Eagle (Cat. No. M0275, Sigma-Aldrich), 10 µl Sodium Bicarbonate 7.5% solution (Gibco), 150 μL Type 1 Bovine Collagen Solution (#5005, PureCol^®^, Advanced BioMatrix, Sweden), and 90 µl cell suspension were sequentially mixed and gently homogenized.

Hydrogels contained either 2*10^5^ fibroblasts alone or a mix of 2*10^5^ fibroblasts with 1*10^6^ PBMCs, monocytes or PBMC depleted from CD14^+^ cells. The hydrogels containing M1 or M2-like macrophages had 5*10^5^ cells together with 2*10^5^ fibroblasts. Of the collagen-cell mixture, 250 µl was used per well in 48-well plates. Hydrogels were incubated under standard culture conditions (37 °C, 5% CO_2_) until solidification for at least 1 hour. Finally, 750 µl of RPMI 1640 GlutaMAX^TM^ (Gibco) supplemented with 100 U/ml penicillin, 100 mg/ml streptomycin, 100 mg/L pyruvate, and 10% Human Pooled Serum (HPS) were added and the hydrogels put back in the culture incubator. In another experimental setting, hydrogels containing only fibroblasts were exposed to M1 or M2-like macrophage culture supernatant. For this, after hydrogels’ solidification, 250 μL of M1 or M2-like culture supernatant was added to each well. One day after (24 hours), 500 μL RPMI medium was added.

In all experiments, three technical replicates were used. Spontaneous contraction of the hydrogel was macroscopically evaluated over time by scanning plates on a standard office flatbed scanner with 600 dpi resolution. Hydrogel contraction was quantified by measuring the area of the hydrogels using Fiji ImageJ software and calculated as follows:

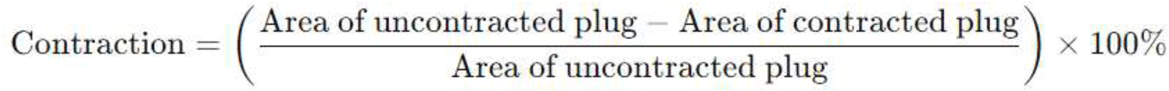

### Drugs and reagents

Tofacitinib (1 µM) (LC Laboratories, USA), SB-505124 (5 µM) (Sigma-Aldrich), IKK-16 (1,5 mM) (MedChemExpress, USA) and TAK-242 (1 mM) (R&D Systems) were diluted into culture medium and subsequently added to the hydrogels after solidification. To analyze the role of soluble mediators possibly released by M1 or M2-like macrophages in myofibroblast contraction, anti-IL-6 (Cat. No. 501101, BioLegend, USA) (1 µg/ml) and anti-TGF-β (#MAB1835, R&D Systems) (100 ng/ml) neutralizing antibodies were used. An IgG isotype (Cat. No. 401501, BioLegend) antibody was used as a control. M1 or M2-like macrophage supernatant was incubated for 1 hour with the respective neutralizing antibodies and then added to the hydrogels.

### Flow cytometry

Collagen hydrogels were enzymatically digested (for 1 hour on a roller at 37°C) using a mixture of collagenase D (1:50) (Roche, Switzerland) and DNAse (1:100) (Roche) diluted in PBS. Reaction was stopped by adding cold PBS. The hydrogels were mechanically disrupted by gently pipetting up and down, then centrifuged at 300 × g for 5 minutes. After centrifugation, supernatant was discarded and the obtained single-cell suspension was washed twice with PBS, followed by isolation of the immune cell fraction using CD45^+^ magnetic MicroBeads (130-045-801, Miltenyi Biotec). The isolated CD45^+^ portion was used to access cells’ phenotype by flow cytometry.

Briefly, leukocytes isolated from the digested hydrogels were first stained for cell viability with ViaKrome 808 (1.5:1000 in PBS) (Beckman Coulter, USA) for 30 minutes at 4°C in the dark. Then, samples were washed and incubated for 20 minutes at room temperature (protected from light) with fluorescently labeled extracellular antibodies. Subsequently, intracellular staining was performed using the Cyto-Fast™ Fix/Perm Buffer Set (BioLegend) following the manufacturer’s instructions. Cells were acquired in a Cytoflex LX 21-color flow cytometer (Beckman Coulter). Data analyses were conducted using Kaluza software version 2.1.3 (Beckman Coulter). A list of all extra-and intracellular antibodies used is provided in **Supplementary Table 1**.

### Immunohistochemistry

Collagen hydrogels were collected and fixed in a 10% buffered formalin solution, followed by embedding in paraffin. Five-micrometer (μm) thick paraffin sections were cut from formalin-fixed blocks and added to glass slides. Identified samples were then deparaffinized with xylol wash and rehydrated with ethanol. Afterward, the material passed by an antigen retrieval process using 10 mM citrate (pH 6.0) at room temperature. Endogenous peroxidase activity was inhibited by incubating the samples with 3% H_2_O_2_ in PBS for 15 min, and subsequently, non-specific binding was blocked using 20% Normal Goat Serum (NGS)/PBS or horse serum. Sections were then incubated with primary rabbit –anti-human alpha-smooth muscle actin (α-SMA) (1:200) (ab5694, Abcam, UK), – fibroblast activation protein (FAP) (1:100) (ab207178, Abcam), – Phospho-SMAD2 (1:100) (#3108L, CellSignaling, USA), – Phospho-Stat3 (1:50) (#9145, CellSignaling), or mouse anti-human – CD68 (#MCA1815, BioRad, USA) (1:200) antibodies, for 1 hour at room temperature. Next, tissues were incubated with corresponding secondary antibodies: goat anti-rabbit biotinylated IgG antibody (1:400) (#BA-1000, VectorLabs, USA) or horse anti-mouse biotinylated IgG antibody (1:200) (#BA-2001, VectorLabs) and then labeled with the avidin-biotin complex (1:100) (VECTASTAIN, Thermo-Fisher). Finally, reaction was revealed using 3,3’-diaminobenzidine (bright DAB, Immunologic). All slides were counterstained with hematoxylin and mounted with a cover slip (Permount, Thermo-Fischer, USA).

### RNA isolation and quantitative real-time PCR

Total RNA was extracted from the CD45^−^ cell fraction (fibroblasts) using the RNeasy Mini Kit (Qiagen, Germany), following the manufacturer’s instructions. RNA concentrations were quantified with a Nanodrop photo spectrometer (Thermo Scientific, USA) and any residual DNA was removed by treating samples with DNAse I (Life Technologies, EUA). Subsequently, RNA was reverse-converted into cDNA in a single step reverse transcription PCR, using oligo dT primer and 200U M-MLV Reverse transcriptase (All Life Technologies, USA), in a thermocycler. Gene expression was evaluated on the produced cDNA samples using SYBR Green Master Mix (Applied Biosystems, USA) and 0.2 mM of validated specific primers (Biolegio, Nijmegen, the Netherlands) in a quantitative real-time polymerase chain reaction (qPCR). Primers sequences are provided in **Supplementary Table 2**. All reactions were carried out in triplicate, using a StepOnePlus Real-Time PCR System (Applied Biosystems). Reference genes used were GAPDH and RPS27A. The relative gene expression (−ΔCt) was calculated using the average values of the reference genes.

### Reporter Luciferase Assay

Reporter constructs cloned in the same human primary skin dermal fibroblasts were obtained from (18). Report activity known to induce relevant signaling pathways on fibroblasts, such as Sis-inducible element (SIE), SMAD binding element (SBE), cyclic AMP response element (CRE), nuclear factor of activated T-cells 5 (NFAT-5), interferon-stimulated response element (ISRE), and nuclear factor-κB response element (NF-κB), were tested (**Supplementary Table 3**). Hydrogels containing monocytes and the different fibroblast constructs were prepared using the previously mentioned cell ratio (5:1). The co-culture was incubated for 24 hours and then cells were lysed using Cell Culture Lysis Reagent (Promega, USA). Next, Nano-Glo luciferase reagent (Promega) was added according to the manufacturer’s protocol. Fluorescence was measured at 590nm with a Clariostar (BMG Labtech, Germany). An appropriate positive control, known to activate each fibroblast construct, was used.

### Statistical analysis

Statistical analyses were performed with GraphPad Prism 10 (10.0.3 version; GraphPad Software Inc., USA). Depending on data normality, either Student’s t-test, Wilcoxon test, one-way ANOVA, or Kruskal–Wallis test was performed. The specific statistical test used for each analysis is indicated in the respective figure legends. Differences were considered significant with a p-value of less than 5% (p < 0.05).

## DATA AVAILABILITY

### Supplementary Information

The online version contains supplementary material. Correspondence and requests for materials should be addressed to A.P.M.v.C.

## ACKNOWLEDGEMENTS

The authors would like to thank the funding provided by public Brazilian grant from the São Paulo Research Foundation (FAPESP), Grant number: 2023/04897-7.

## AUTHOR CONTRIBUTIONS

Conception and design: D.C.Z.S., M.I.K. and A.P.M.v.C. Collection and acquisition of data: D.C.Z.S., N.J.T.v.K., T.I.P., B.W. and E.L.V. Analysis and interpretation of data: D.C.Z.S., N.J.T.v.K., T.I.P., D.N.D., M.H.J.v.d.B., M.I.K. and A.P.M.v.C. Drafting of the manuscript: D.C.Z.S., M.I.K. and A.P.M.v.C. Critical revision: D.C.Z.S., N.J.T.v.K., T.I.P., D.N.D., B.W., E.L.V., M.H.J.v.d.B., M.I.K. and A.P.M.v.C. Final approval of the article: D.C.Z.S., N.J.T.v.K., T.I.P., D.N.D., B.W., E.L.V., M.H.J.v.d.B., M.I.K. and A.P.M.v.C. All authors have read and agreed to the submitted version of the manuscript.

## ADDITIONAL INFORMATION

The authors declare no competing interests.

