## Supplementary figures and tables for "Monocytes Strongly Induce (MYO)Fibroblast Contraction in a New 3D Skin Model to Understand the Inflammation-Fibrosis Axis in Systemic Sclerosis": Supp. figures legends.docx

**Supplemental Figures**

**Supplemental figure 1. Additional transcription factor–responsive reporter constructs tested in fibroblast-monocyte co-culture**. Dermal fibroblasts expressing ISRE (**A**), NFAT-5 (**B**), or CRE (**C**) luciferase reporter constructs were cultured with monocytes in collagen hydrogels for 24 hours. Luminescence was measured as a readout of reporter activity. Fibroblast-only hydrogels were included as controls. Each dot represents the mean of three technical replicates from one PBMC donor (n = 3). Statistical analysis was performed using Student’s t-test (**p < 0.01). ISRE: Interferon-stimulated response element (ISRE); NFAT-5: Nuclear factor of activated T-cells 5, and CRE: Cyclic AMP response element.

**Supplemental figure 2. Blocking the TLR4 signaling did not affect (myo)fibroblasts contraction.** Co-cultured collagen hydrogels were treated with TAK-242 (1 µM) or vehicle (DMSO), and contraction was measured after 72 hours. (**A**) Quantification of hydrogel contraction showing no significant differences between control and TAK-242–treated conditions, indicating that TLR-4 inhibition had no impact to halt myofibroblasts contraction. Each symbol represents the mean of three technical replicates from one PBMC donor (n = 5). Statistical analysis was performed using Student’s t-test. (B) Representative scanned images of co-cultured hydrogels treated with TAK-242.

**Supplemental figure 3. Flow cytometry gating strategy for monocytes-containing hydrogel plugs.** Collagen hydrogels co-cultured with fibroblasts and monocytes were enzymatically digested to obtain a single-cell solution, followed by MACS-CD45^+^ isolation and flow cytometric analyses. Gating was performed sequentially on single, live cells, total leukocytes (CD45^+^). Within the leukocyte population, macrophages were identified as CD45⁺CD68⁺ cells. Macrophages were further subdivided into M1-like (CD163⁻HLA-DR⁺CD86⁺) and M2-like (CD163⁺CD206⁺) subsets.

**Supplemental figure 4. Interaction with fibroblasts drives monocytes toward M1-M2 mixed macrophage polarization phenotype**. Representative flow cytometry plots of macrophage subsets isolated from collagen hydrogels co-cultured with fibroblasts and monocytes. (**A**) M1-like macrophages (CD163⁻) were gated for CD86⁺HLA-DR⁺ but also exhibited detectable CD206 expression. (**B**) M2-like macrophages (CD163⁺) expressed canonical M2 markers (CD206⁺) while also showing M1-associated markers (CD86⁺HLA-DR⁺).

**Supplemental figure 5**. **Fibroblasts reporter constructs co-cultured with M1- or M2-like macrophages.** Dermal fibroblasts expressing NF-κB (**A**), ISRE (**B**), NFAT-5 (**C**), or CRE (**D**) reporter constructs were cultured with either M1- or M2-like macrophages, and luciferase activity was measured for respective signaling reporters. Fibroblast-only hydrogels were included as controls. Each shape represents the average of 3 technical replicates of 1 PBMC donor, n = 3. Statistical analyses were performed using Student’s t-test (****p<0.0001). NF-κB: Nuclear factor-κB response element; ISRE: Interferon-stimulated response element (ISRE); NFAT-5: Nuclear factor of activated T-cells 5, and CRE: Cyclic AMP response element.
