## Supplementary figures and tables for "Monocytes Strongly Induce (MYO)Fibroblast Contraction in a New 3D Skin Model to Understand the Inflammation-Fibrosis Axis in Systemic Sclerosis": Supplementary Tables.docx

**Supplementary Table 1: List of antibodies used for extra- and intracellular staining**

| **Antigen** | **Clone** | **Dilution** | **Fluorochrome** | **Supplier** |
| --- | --- | --- | --- | --- |
| CD45 | HI30 | 1:100 | APC-Fire 750 | BioLegend |
| CD86 | BU63 | 1:200 | PerCP/Cy5.5 | BioLegend |
| HLA-DR | L243 | 1:50 | BV421 | BioLegend |
| CD163 | GH1/61 | 1:200 | BV711 | BioLegend |
| CD206 | 15-2 | 1:200 | FITC | BioLegend |
| CD68* | Y1/82A | 1:100 | PE | BioLegend |

*CD68 was used for only intracellular staining

**Supplementary Table 2: List of primer sequences used for qPCR**

| **Gene** | **Forward primer 5’ – 3’** | **Reverse primer 5’ – 3’** |
| --- | --- | --- |
| GAPDH | ATCTTCTTTTGCGTCGCCAG | TTCCCCATGGTGTCTGAGC |
| RPS27A | TGGCTGTCCTGAAATATTATAAGGT | CCCCAGCACCACATTCATCA |
| PDPN | GGTGCAATCATCGTTGTGGTTA | TTCAGCTCTTTAGGGCGAGTAC |
| COL1A1 | AGATCGAGAACATCCGGAG | AGTACTCTCCACTCTTCCAG |
| IL6 | AGCCCACCGGGAACGA | GGACCGAAGGCGCTTGT |
| FN1 EDA | TTCAGACTGCAGTAACCAACAT | GGTCACCCTGTACCTGGAAAC |
| COL3A1 | CCTGGAATCTGTGAATCATGCC | TGCGAGTCCTCCTACTGCTA |
| ACTA2 | CTGACCCTGAAGTACCCGATA | GAGTGGTGCCAGATCTTTTCC |
| HLA-DR | CCCTGCAGCACCACAAC | GGAACCACCTGACTTCAATGC |
| PLOD2 | AAGACTCCCCTACTCCGGAAA | AGCAGTGGATAATAGCCTTCCAA |

**Supplementary Table 3: Luciferase constructs for transcription factor (TF) activity utilized, each recognized for activating key signaling pathways in fibroblasts**

| Pathway | TF(s) | TF Element (Abbreviation Construct) | Positive Control | Reference |
| --- | --- | --- | --- | --- |
| NFκB | NFκB:p65 | Nuclear factor κ B  response element (NFκB) | IL-1β  (1 ng/mL) | OHMORI; HAMILTON, 1995 |
| cAMP/PKA | CREB | Cyclic AMP  response element (CRE) | Forskolin (10 µM) | TAN et al., 1994 |
| TGF-β | SMAD3:SMAD4 | SMAD binding  element (SBE) | TGF-β1  (1 ng/mL) | REISDORF  et al., 2001 |
| INF-α | STAT1:STAT2 | Interferon-stimulated  response element (ISRE) | IFN-α  (100 ng/mL) | OHMORI; HAMILTON, 1995 |
| IL-6 | STAT3:STAT3 | Sis-inducible element (SIE) | IL-6  (10 ng/mL) | CHAKRABORTY et al., 2017 |
| Calcium/calcineurin:  hyperosmotic signaling | NFAT5 | Nuclear factor of activated  T-cells 5 | +100 milliosmole | HERBELET  et al., 2018 |
