## Supplementary figures and images for "Monocytes Strongly Induce (MYO)Fibroblast Contraction in a New 3D Skin Model to Understand the Inflammation-Fibrosis Axis in Systemic Sclerosis"

### Supp Fig 1.png

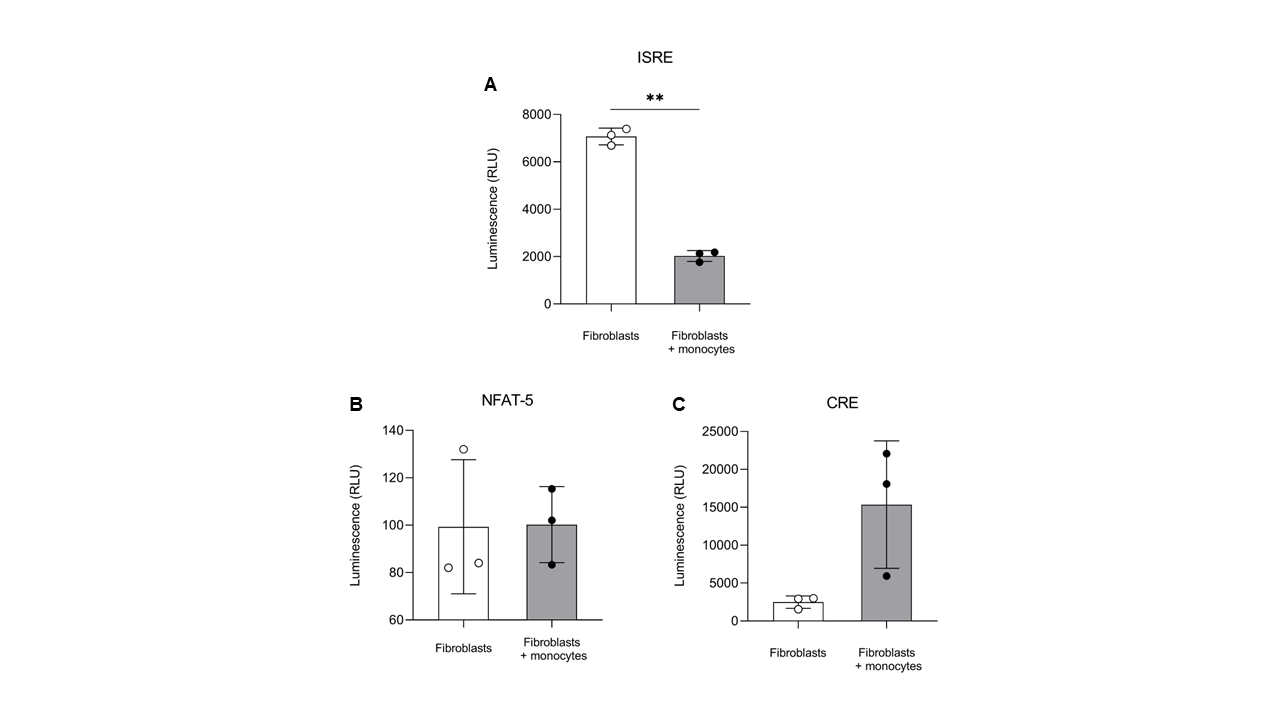

### Supp Fig 2.png

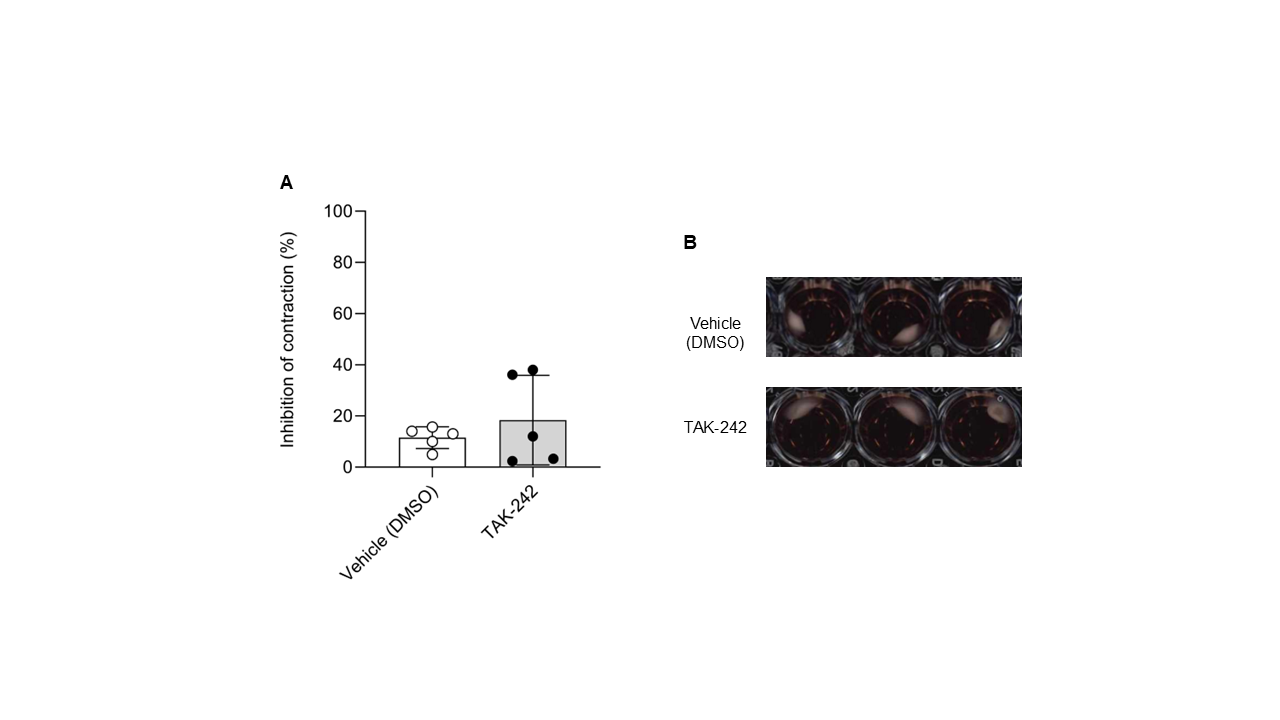

### Supp Fig 3.png

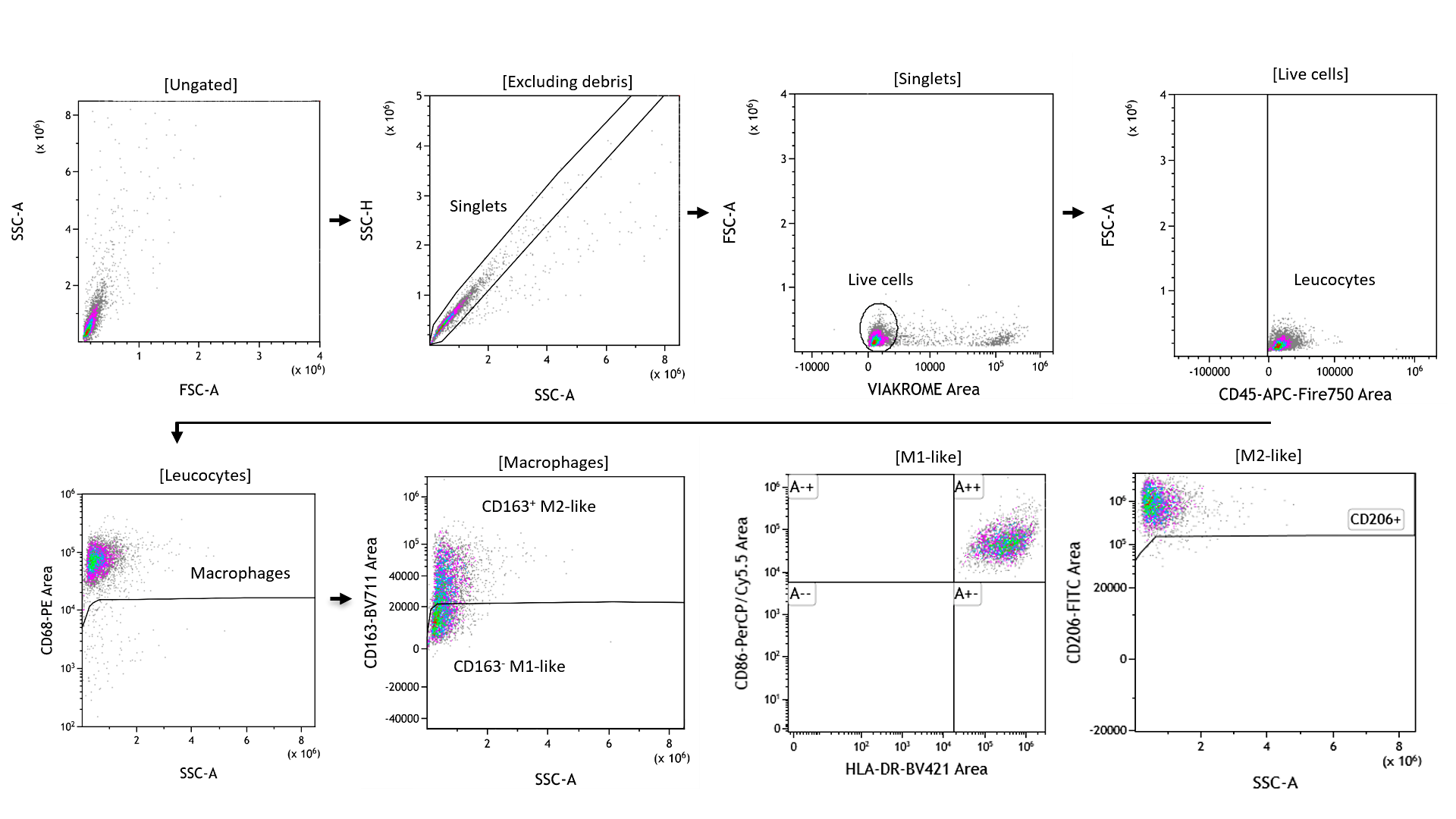

### Supp Fig 4.png

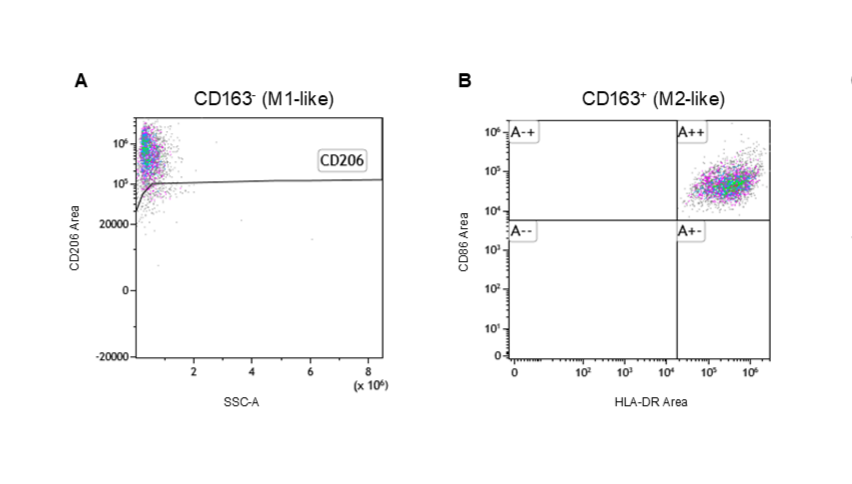

### Supp Fig 5.png

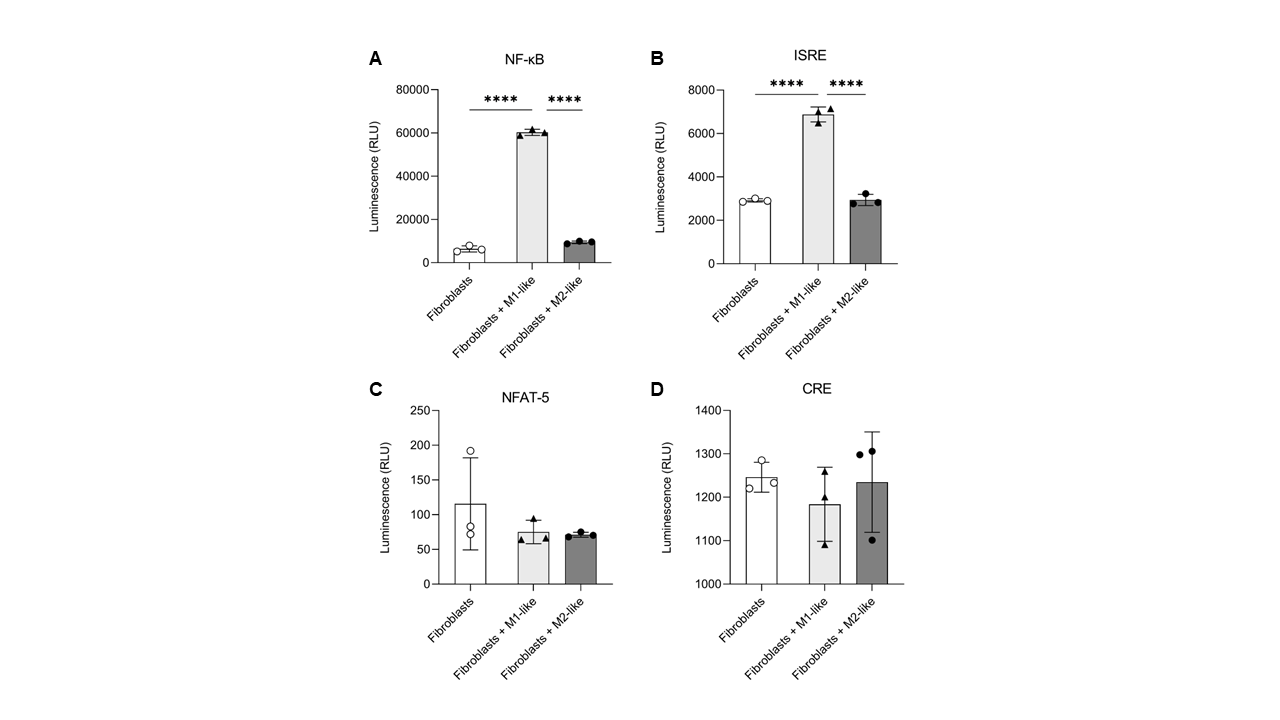
